# Extraction and analysis of methylation features from Pacific Biosciences SMRT reads using MeStudio

**DOI:** 10.1101/2022.03.23.485463

**Authors:** Christopher Riccardi, Iacopo Passeri, Lisa Cangioli, Camilla Fagorzi, Alessio Mengoni, Marco Fondi

## Abstract

**Motivation:** DNA methylation is the most relevant epigenetic information, present in eukaryotes and prokaryotes, and is related to several biological phenomena, from cellular differentiation to control of gene flow, pathogenesis and virulence. The widespread use of third-generation sequencing technologies allows direct and easy detection of genome-wide methylation profiles, offering increasing opportunities to understand and exploit the epigenomics landscape.

**Results:** We introduce MeStudio, a pipeline which allows to analyse and combine genome-wide methylation profiles with genomic features. Outputs report the presence of DNA methylation in coding sequences, noncoding sequences, intergenic sequences, and sequences upstream to CDS. We show the usage and performances of MeStudio on a set of single-molecule real time sequencing outputs from the bacterial species *Sinorhizobium meliloti*.

**Availability and Implementation:** MeStudio is written in Python, Bash and C and is freely available under an open source GPLv3 license at https://github.com/combogenomics/MeStudio

**Supplementary information:** Supplementary data are available at *Bioinformatics* online.

## 1 Introduction

DNA methylation has been shown to be pivotal in the control of several biological phenomena in eukaryotes and prokaryotes (Jones, 2012). Third-generation sequencing technologies, namely single molecule real-time (SMRT) (Flusberg *et al*., 2010; Fang *et al*., 2012) and nanopore ONT (Clarke *et al*., 2009; Simpson *et al*., 2017) sequencing allow to directly identify the most commonly methylated bases (Gouil and Keniry, 2019; Sánchez-Romero and Casadesús, 2020; Rand *et al*., 2017). These methods are rapidly increasing the number of studies on genome-wide DNA methylation, especially in prokaryotes, where the compact size of genomes allows the generation of whole-genome methylome with relative ease. Consequently, the interest toward computational pipelines which can dig into DNA methylation features in a genome-wide manner is growing. Several tools have been developed for the analysis of DNA methylation profiles deriving from bisulphite sequencing and microarrays (e.g. (Müller *et al*., 2019; Teng *et al*., 2020; Hillary and Marioni, 2021; Aryee *et al*., 2014; Bock *et al*., 2005)), for a recent benchmarking see (Nunn *et al*., 2021)). Recently, three packages have been released (Su *et al*., 2021; Leger, 2020; De Coster *et al*., 2020), which allow to visualize methylation profiles from SMRT or ONT sequencing data. However, to the best of our knowledge, no specific pipeline has been developed for extracting DNA methylation information and allowing a direct quantification/comparison of the position of methylated sites with respect to genome-derived features, such as coding and noncoding sequences. Here we present MeStudio, a pipeline for SMRT sequencing data integration and visualization. MeStudio combines methylation data with genome sequence and annotation to facilitate the extraction of biological information from DNA methylation profiles and to visualize the results of these analyses. We show the usage of MeStudio on a set of SMRT outputs from the bacterial species *Sinorhizobium meliloti*.

## 2 Design and implementation

MeStudio connects several tools that can be run individually or as part of a pipeline. The required input data consist in only three files: i) a FASTA file containing the genome sequence, ii) a genomic annotation file in GFF3 format and iii) another GFF3 containing the methylated nucleotide positions. The latter is automatically generated from the output of the SMRTlink software of Pacific Biosciences DNA sequencers. MeStudio outputs text files containing the information of the methylated sites, according to their corresponding genomic region (see below). A workflow is provided in Figure 1.

**Figure 1:**
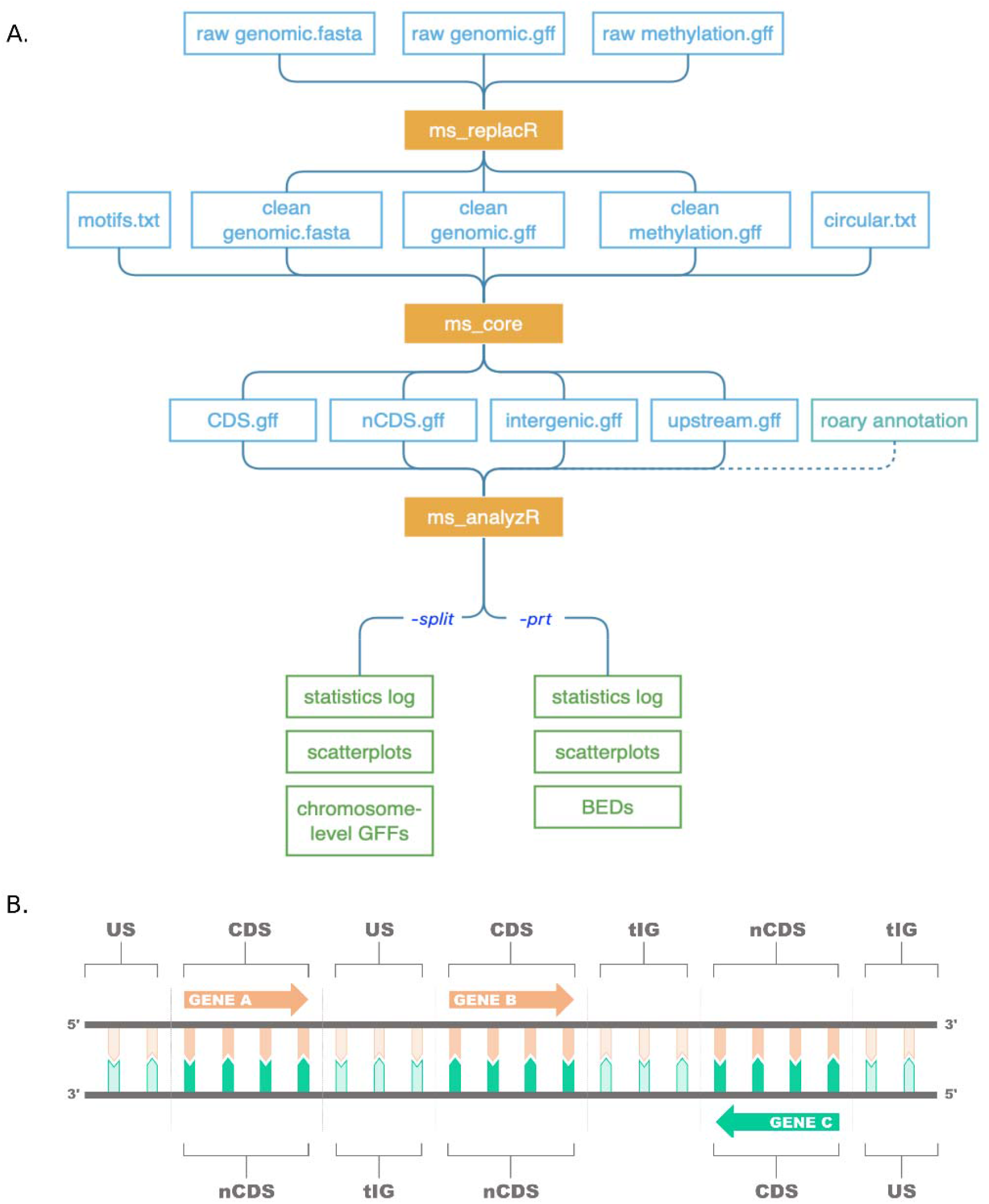
MeStudio overview A) Workflow. Each blue block represents input files. The yellow blocks indicate the scripts. The green boxes indicate final results files. B) Graphical representation of CDS, nCDS, tIG and US. See text for details.

### 2.1 Pre-processing

To run MeStudio, a pre-processing python script named *ms_replacR* has been implemented to produce consistent formatting on the sequence identifiers from the genomic annotation, sequencer-produced modified base calls, and the genomic sequence file. More details are provided in the MeStudio manual on GitHub.

### 2.2 Core-processing

The processing of the input files is handled by five executables which we refer to as MeStudio core. These components match the nucleotide motifs to the genomic sequence and map them to the corresponding category, which are extracted from the annotation file. Categories are defined as follows: i) protein-coding gene with accordant (sense) strand (CDS), ii) discordant (antisense) strand (nCDS), iii) regions that fall between annotated genes (true intergenic, tIG), iv) regions upstream to the reading frame of a gene, with accordant strand (US) (Figure 1B). The current implementation uses a naive matching algorithm to map motif sequences to the reference genome. During the naive matching phase, each replicon or chromosome gets loaded into memory in the form of an array of chars (or null terminated C-string) one at a time as both strands are scanned for the presence of the motif sequences, which can hold ambiguity characters. The resulting binary files are then processed by another executable that is called for the task at hand. MeStudio core crosses methylated bases positions relative to the reference sequence start with the previously described features, producing GFF3 files that serve as input for the final analysis stage. This is an expensive part of the pipeline in which multiple nested for loops and calculations are performed. Integrating one motif on a four-contigs genome (6,973,268 bp, 23,433 GANTC motif matches) took 0m27.116s on a single AMD Opteron 6380 processor (2.5GHz).

### 2.3 Post-processing

MeStudio implements a post-processing python script named *ms_analyzR* which takes MeStudio core output as input. In addition, to integrate comparative genomic analyses a “gene_presence_abscence.csv” file produced by Roary (Page *et al*., 2015) can be used to define the methylation level and patterns of core and dispensable genome fractions, as well as to annotate the genes-coded proteins. The script logs to the standard output some statistics and information: in particular, *ms_analyzR* logs the total number of genes found for each category (CDS, nCDS, tIG, US). Additionally, methylation data are shown, such as i) total number of methylated sites, ii) total number of methylated genes, iii) the ID of the most methylated gene (geneID) and, iv) the product of that gene. Notably, integrating data from Roary is functional to characterize the geneID associated with the name of the protein (as annotated by Prokka) as part of the core or dispensable genome. All the information is saved into a log file, together with plots accounting for the distribution of the methylations (Supplementary Fig. 1). *ms_analyzR* also includes two optional flags named “split” and “prt”. The “split” flag saves into the previously specified output directory the GFFs at “chromosome level” rather than “category level”. Each GFF produced will be characterized not by category (CDS, nCDS, tIG and US) but by chromosomes (or contigs), maintaining the MeStudio core-derived contents and layout unaltered. The “prt” flag produces a BED file for each feature in which is reported: i) the *chrom* column, with the name of each chromosome or contig, ii) start and iii) end of the feature, iv) the name of the geneID found in that interval, v) the number of methylations found for geneID and lastly vi) the protein product of the ID. Information contained in BED files can be readily used to plot the distribution of the methylation density for each feature, making use of the *circlize* R package (https://github.com/jokergoo/circlize) (Supplementary Fig. 2).

## 3 Case study

In order to show the performances of MeStudio, a mock dataset and a recently published SMRT dataset were used (diCenzo *et al*., 2022). Demo files for input and output are available at https://github.com/combogenomics/MeStudio.

