## Supplementary figures and images for "Extraction and analysis of methylation features from Pacific Biosciences SMRT reads using MeStudio"

### Supplementary Figure 1

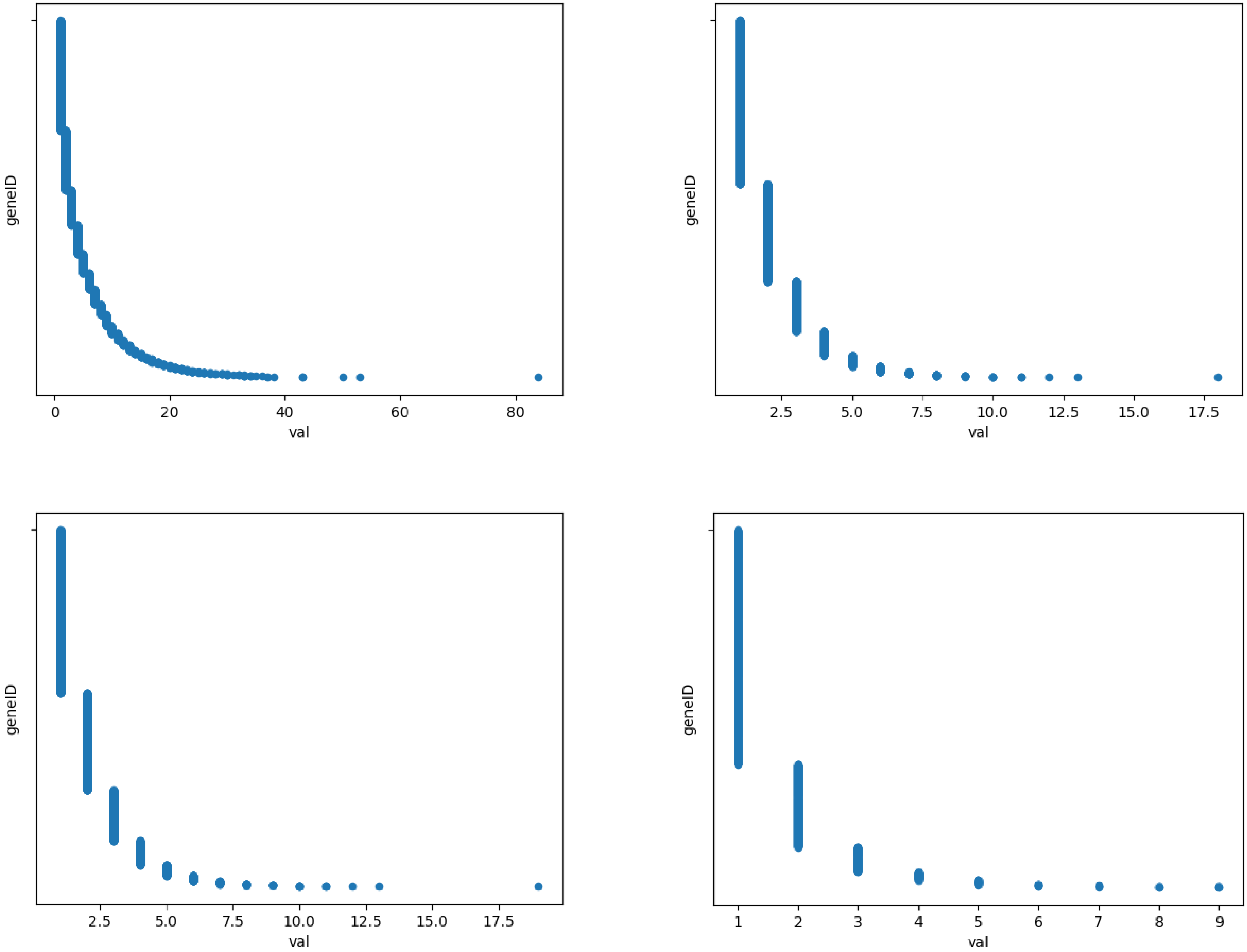

### Supplementary Figure 2

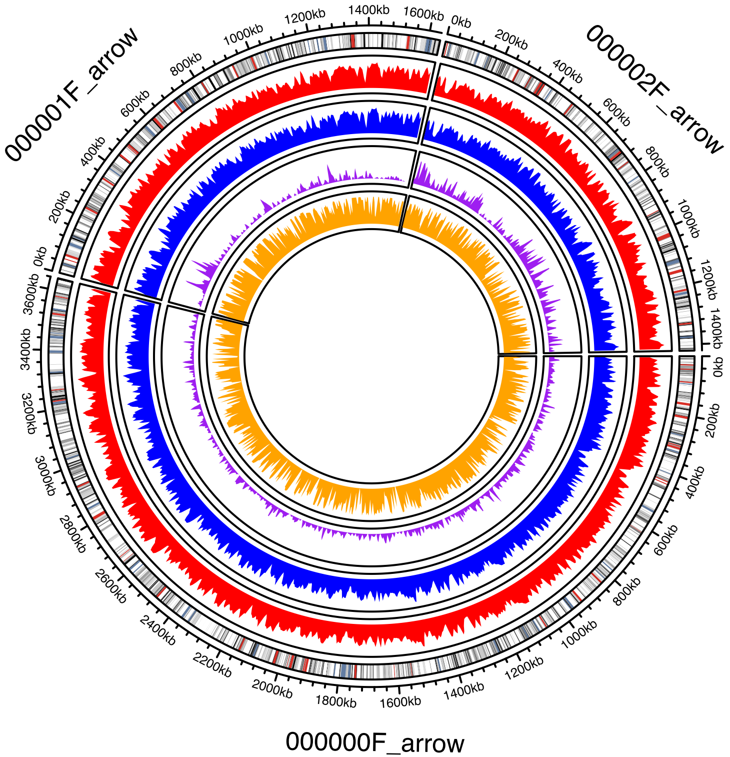
